# Disease specific alterations in the olfactory mucosa of patients with Alzheimer’s disease

**DOI:** 10.1101/2020.11.24.395947

**Authors:** Riikka Lampinen, Mohammad Feroze Fazaludeen, Simone Avesani, Tiit Örd, Elina Penttilä, Juha-Matti Lehtola, Toni Saari, Sanna Hannonen, Liudmila Saveleva, Emma Kaartinen, Francisco Fernandez Acosta, Marcela Cruz-Haces, Heikki Löppönen, Alan Mackay-Sim, Tarja Malm, Minna U Kaikkonen, Anne M Koivisto, Anthony R White, Rosalba Giugno, Sweelin Chew, Katja M Kanninen

## Abstract

Olfactory dysfunction manifests early in several neurodegenerative disorders. Olfaction is orchestrated by olfactory mucosal cells located in the upper nasal cavity. However, it is unclear how this tissue reflects key neurodegenerative features in Alzheimer’s disease. Here we report that Alzheimer’s disease olfactory mucosal cells obtained from live individuals secrete toxic amyloid-beta. We detail cell-type-specific gene expression patterns, unveiling 147 differentially expressed disease-associated genes compared to the cognitively healthy controls, and 5 distinct populations in globose basal cell -, myofibroblast-, and fibroblast/ stromal – like cells *in vitro*. Overall, coordinated alteration of RNA and protein metabolism, inflammatory processes and signal transduction were observed in multiple cell populations, suggesting a key role in pathophysiology. Our results demonstrate the potential of olfactory cell cultures in modelling Alzheimer’s disease advocate their use for diagnostic purposes. Moreover, for the first time we provide single cell data on olfactory mucosa in Alzheimer’s disease for investigating molecular and cellular mechanisms associated with the disease.

## INTRODUCTION

The olfactory mucosa (OM) is critical for the sense of smell and in direct contact with the brain ^1^. Human OM cells can be obtained for *in vitro* studies with a relatively non-invasive biopsy, thus providing a way of gaining access to neural tissue of living individuals ^2^. Anosmia, loss of the sense of smell, has been linked to the early phases of several neurodegenerative diseases including Alzheimer’s disease (AD) ^1,3–5^. AD pathology is typified by extracellular beta-amyloid (Aβ) plaques, intracellular neurofibrillary tangles consisting of hyperphosphorylated tau protein, cell loss, neuroinflammation, and metabolic stress. Young-onset or familial AD is rare and caused by mutations in genes encoding the y-secretase complex members presenilin 1 (PS1) or presenilin 2 (PS2), and in the amyloid precursor protein (APP). However, late-onset AD (LOAD) is the most common disease form and the major risk gene for this is *APOE*, the ε4 allele of which confers a 12-fold increase in disease risk in homozygous carriers ^6^.

Nasal secretions of LOAD patients contain elevated amounts of Aβ and phosphorylated tau ^7–9^, indicative of pathological processes occurring in the nasal cavity. *Post mortem* histological analysis of AD OM have revealed immunostaining for filamentous tau inside neurites and in neuronal soma, resembling intracellular neurofibrillary tangles, in cells close to the OM basal membrane ^10^. Intracellular Aβ and increased Aβ aggregation in the apical surface of the epithelium have also been found in histological sections of OM biopsies derived from living anosmic LOAD patients ^11^. However, little information exists on cell-type-specific, AD-related alterations to OM cells. The only reported cell-type-specific observations thus far are from olfactory neurons in *post mortem* analyses of AD patients ^12,13^, and from OM-derived neuroblast cultures that exhibit elevated oxidative stress and altered APP processing ^14,15^. The cellular content of the biopsied OM of healthy individuals was recently assessed by single-cell RNA sequencing (scRNA-seq) ^16^, but cell-type-specific alterations occurring in AD OM at the single-cell level remain unknown. Human OM cells can be obtained for *in vitro* studies with a relatively non-invasive biopsy, thus providing a way of gaining access to neural tissue of living individuals ^2,16,17^.

In this study, we harvested OM biopsies from cognitively healthy individuals and age-matched patients with LOAD in order to profile AD-linked functional alterations and changes in gene expression at the single-cell level. We observed significantly increased secretion of pathological Aβ from the cultured AD OM cells and 147 differentially expressed genes (DEGs) associated with AD, including cell-type-specific DEGs. These DEGs enriched for pathways including RNA and protein metabolism, inflammatory processes and signal transduction. Together, these results support the utility of the OM as a physiological *in vitro* model of AD and provide a unique cellular-level view of transcriptional alterations associated with AD pathology.

## RESULTS

### Secretion of Aβ_1-42_ is increased in AD OM cells

To test whether OM cells from AD donors exhibit the typical pathological hallmarks of the disease, we first analyzed intracellular and secreted levels of Aβ_1_-_42_, Aβ_1-40_, total tau, and tau phosphorylated at threonine 181 *(P_181_-tau)* in cognitively healthy individuals and patients with AD using ELISA (Fig. 1). A significant increase of secreted Aβ_1-42_ was observed in AD OM cells (18.52 ± 7.590 pg/mg protein, *p*= 0.0366) when compared to cells derived from cognitively healthy individuals. In addition, the ratio of secreted Aβ_1-42_ over Aβ_1-40_ was higher in AD OM cells, in comparison to controls (0.2230 ± 0.06919, *p*=0.0096). No difference was observed in levels of Aβ_1-40_, total tau, or P_181_-tau. Likewise, these proteins remained unaltered in the plasma of AD patients. The levels of secreted Aβ_1-42_ were not found to correlate to the status of the donor’s sense of smell, however, both anosmic and hyposmic individuals were not included in all experimental groups (Supplementary figure 1a). Nevertheless, the OM cells of the AD patients with hyposmia secreted notably increased amounts of Aβ_1-42_ (30.42 ± 13.34 pg/mg protein, p = 0.0847) in comparison to the cognitively healthy individuals with hyposmia. There was no significant difference in the secretion Aβ_1-42_ from OM cells obtained from donors with at least one APOE ε4 allele compared to donors with two APOE ε3 alleles (Supplementary figure 1b).

**Figure 1.**
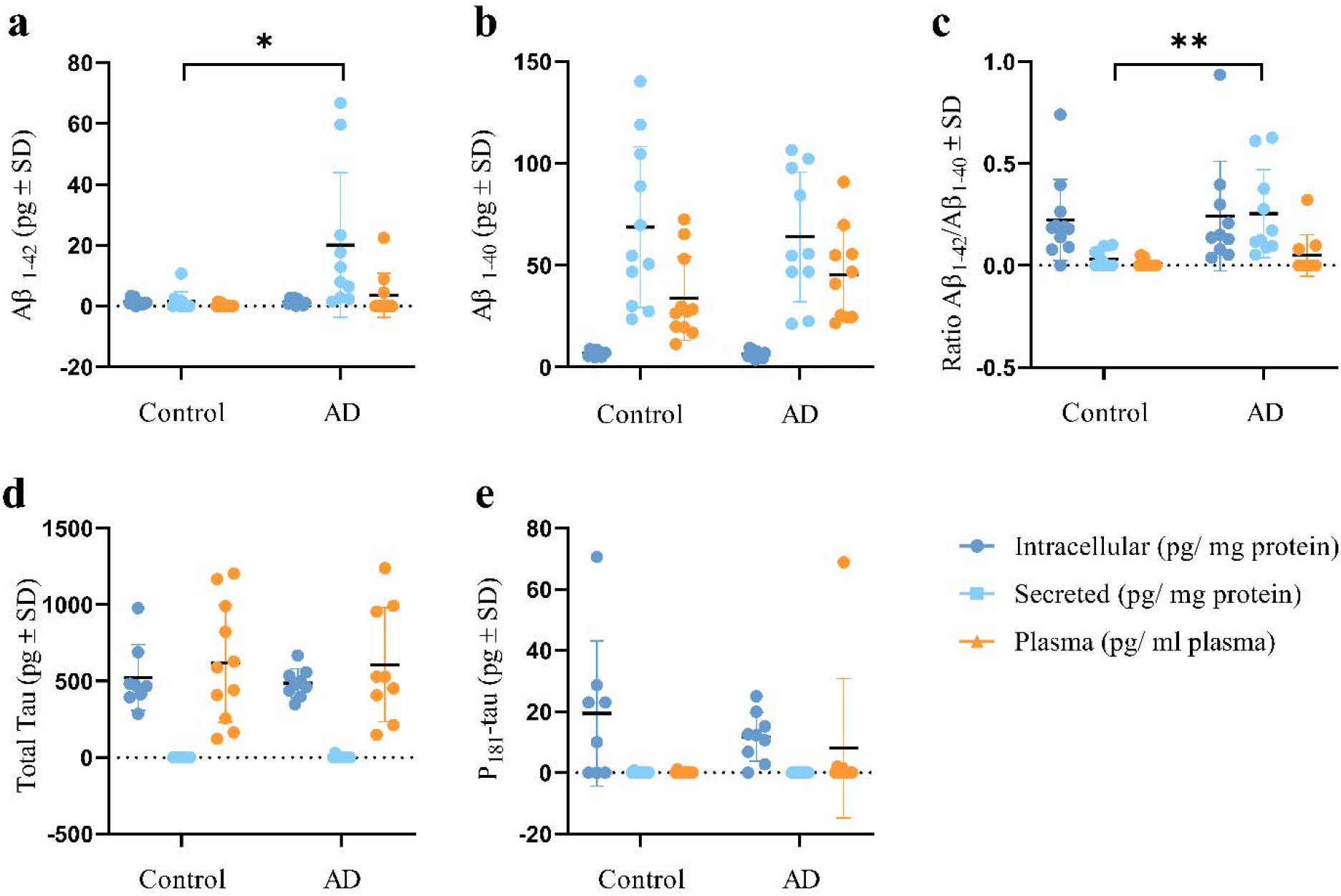
Aβ secretion is increased in AD OM cells. OM cells harvested from biopsies were plated cultured for 7 days prior to collection of culture media and cell lysates for ELISA assays. The results were normalized to the total amount of protein measured from cell lysates. **a)** ELISA assay for Aβ_1-42_. **b)** ELISA assay for Aβ_1-40_. **c)** Ratio of Aβ_1-42_ over Aβ_1-40_. In Aβ ELISA assays N=11 for cognitively healthy controls and N=10 for AD patients. **d)** ELISA assay for total tau. **e)** ELISA assay for P_181_-tau. In tau ELISA assays N=8 for cognitively healthy controls and N=9 for AD patients. Quantification of secreted and intracellular Aβ_1-42_, Aβ_1-40_, tau and P_181_-tau between control and AD OM cells were performed with t test (unpaired, two-tailed, Welch’s correction). **a** * p◻<◻0.05. **c** ** p < 0.01. For all graphs data is presented as mean ± SD and calculated the statistics as difference in means ± SEM.

### Cytokine secretion is not altered in AD OM cells

To profile the innate immunity of OM cells, we next used the CBA array to analyze the levels of 5 cytokines in OM cells and plasma of cognitively healthy control subjects and AD patients. Secreted or intracellular cytokine levels of AD OM cells were not altered in comparison to OM cells from cognitively healthy subjects (Figure 2). However, the plasma samples of AD patients contained significantly increased amounts of tumor necrosis factor (TNF) compared to plasma collected from cognitively normal individuals (1.324 ± 0.2277 pg/ml plasma, *p* < 0.0001) (Fig. 2f).

**Figure 2.**
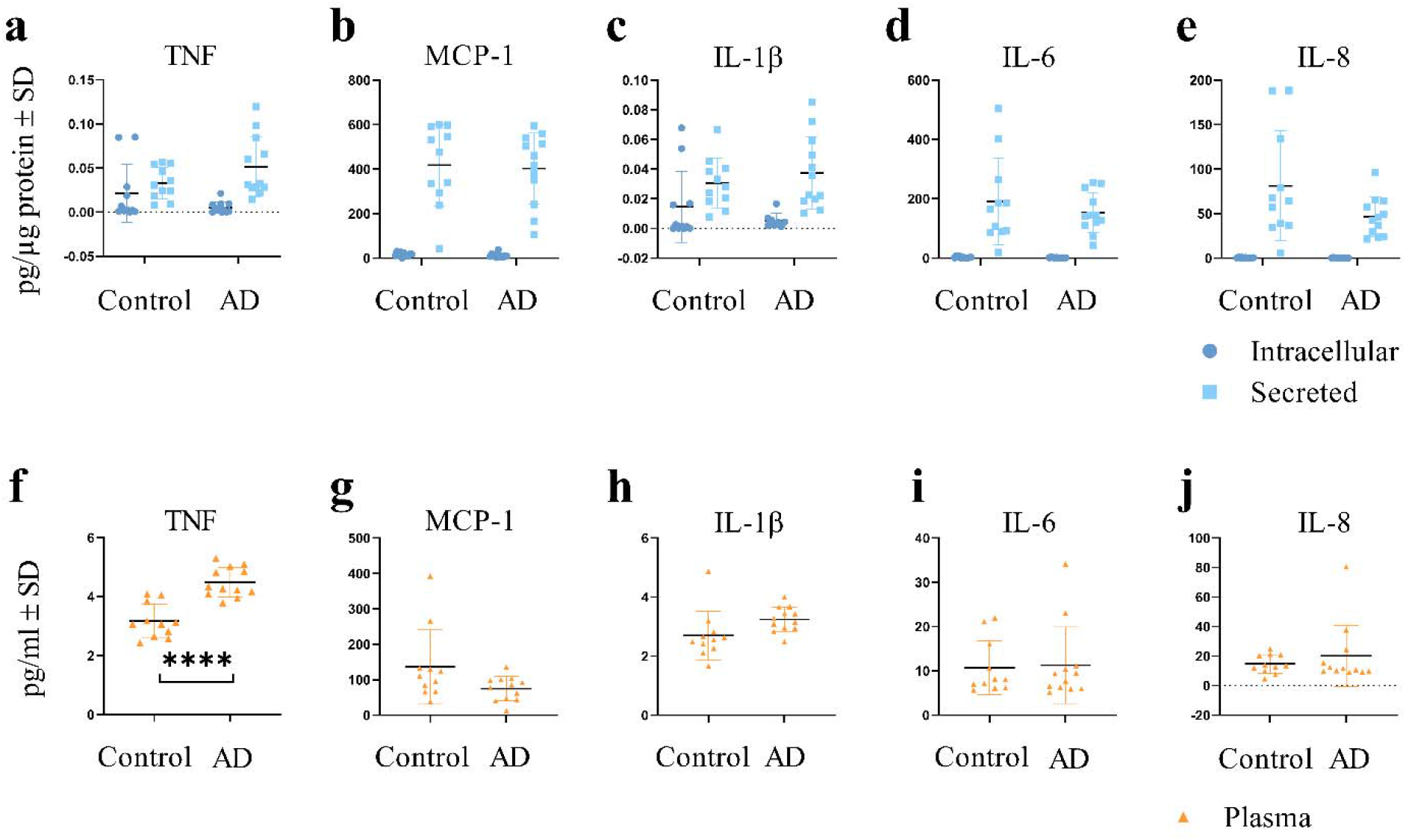
Cytokine levels are unaltered in AD OM cells. OM cells harvested from biopsies were plated and cultured for 7 days prior to collection of culture media and cell lysates for the CBA array. The results were normalized to the total amount of protein measured from cell lysates. **a-e)** Levels of intracellular and secreted cytokines in OM cells. N = 11 for cognitively healthy controls and N = 12 for AD patients. **f-j)** Levels of cytokines in plasma samples. N = 11 for cognitively healthy controls and N = 12 for AD patients. Quantification of secreted, intracellular or plasma cytokines between control and AD cells was performed with t test (unpaired, two-tailed, Welch’s correction). **** p < 0.0001. For all graphs data is presented as mean ± SD and calculated the statistics as difference in means ± SEM. TNF: tumor necrosis factor, MCP-1: monocyte chemoattractant protein 1, IL: interleukin.

### Cellular diversity of the human OM in health and AD

To investigate the cell type diversity and AD-associated cellular changes in the OM, we performed scRNA-seq of OM cultures derived from cognitively healthy controls and patients with AD (Fig. 3a). To reduce variability and exclude the possible effect of *APOE* ε4 allele on the observed results, OM cell lines from female donors with an *APOE* ε3/3 genotype (age average 72.4) were used for scRNA-seq.

**Figure 3.**
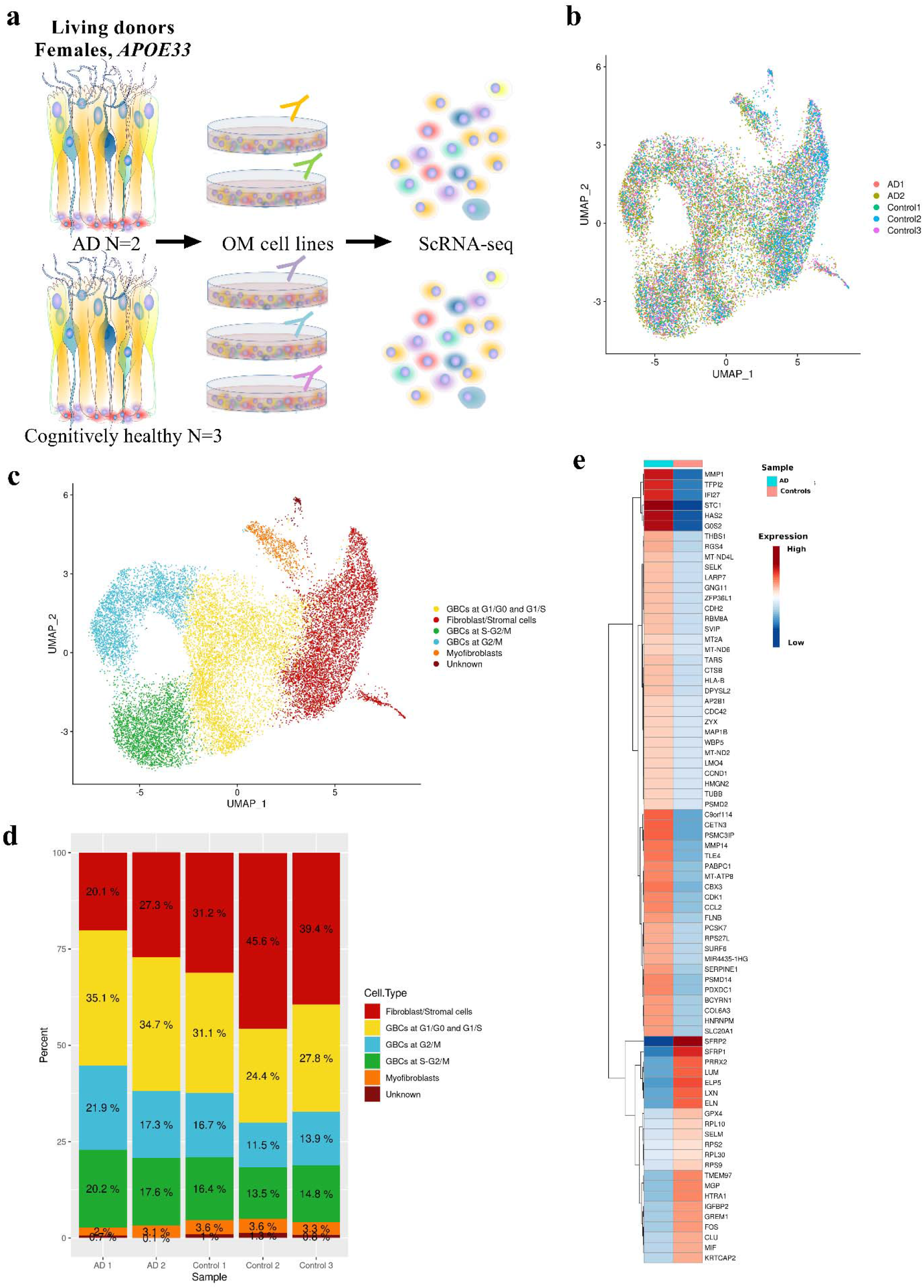
scRNA-seq data revealed AD-specific alterations in OM cells. **a)** Schematic illustration of the sample material and scRNA-seq workflow. **b)** UMAP visualization of the clustering of single cells and showing by colors the cells derived from individual cognitively healthy donors or patients with AD. **c)** Annotated cell types in samples derived from cognitively healthy individuals and patients with AD. **d)** Proportion (%) of cells derived from each donor per cell type. **e)** Heatmap depicting a subset of the AD-related genes differentially expressed between AD and control libraries. The data is presented as the logarithm of the average expression in AD and control cells of the most differentially expressed genes (avg_logFC < −0.3, avg_logFC > 0.3).

We sequenced 10,816 live cells for the control library and 12,582 live cells for the AD library. The median number of genes per cell in the control and AD libraries were 4,066 and 3,336 respectively. After quality-control filtering 8559 cells for controls and 10086 for AD were obtained (Fig. 3b, suppl fig 2). Single cell transcriptomes were clustered in uniform manifold approximation and projection (UMAP) to assess cellular heterogeneity with the Seurat clustering combined with EnrichR and HumanBase tools, and data obtained from Durante et al. ^16^. After data integration, six clusters were identified, consisting of fibroblast/stromal -like cell, globose basal cell (GBC) -like cells at different cell cycle phases and under differentiation, myofibroblast-like cells, and a small cluster of cells annotated as unknown (Fig. 3b-c, suppl. table 1, suppl. fig. 3). Figure 3d and supplementary table 2 show the proportion of cell types present in the OM cultures of each donor. By proportion, the largest number of cells in the OM cultures were annotated as GBC-like and fibroblast/ stromal cell -like cells. In AD the proportion of all GBC-like cells was slightly higher, approximately 73 %, from the total number of cells/ donor, compared to OM cells derived from cognitively healthy individuals were the proportion of all GBC-like cells ranged from 49.4% to 64.2%. A small proportion of cells was annotated as myofibroblast-like cells in both libraries. A maximum of only 1.3% of cells / donor remained unknown (Fig. 3d, suppl. table 2). These cell type categories were used to characterize AD-related gene expression perturbations occurring in the OM on the level of single cells.

### Differential expression analysis reveals AD-related alterations of gene expression in the OM

The single cell transcriptomic analysis of OM cells revealed a total of 147 AD-associated genes differentially expressed between the OM cell libraries of cognitively healthy control individuals and patients with AD (Suppl. table 3). 90 of these DEGs were up-regulated and 57 down-regulated in the AD OM cells. Figure 3e illustrates a subset of the most differentially expressed AD-associated genes between the OM libraries. *STC1*, *TFPI2*, *MMP1*, *CBX3*, *PABPC1*, and *MT-ATP8* were the top six up-regulated DEGs and *SFRP2*, *MGP*, *SFRP1*, *IGFBP2*, *HTRA1*, and *PRRX2* formed the top six most down-regulated DEGs. Given the increased secretion of Aβ_1-42_ from the AD OM cells, we next investigated whether genes linked to APP processing were differentially expressed in the AD OM cells. Gene list analysis of the 90 AD-associated genes up-regulated in the AD OM cells with PANTHER (Protein Analysis Through Evolutionary Relationships, ^18,19^) revealed three genes (*PCSK7, MMP1, MMP14*) annotated to the Alzheimer’s disease-presenilin pathway (PANTHER pathway P00004). In addition, *PCSK7* annotated to the Alzheimer’s disease-amyloid secretase pathway (PANTHER pathway P00003).

### Analysis of individual cell types reveals distinctive AD-associated genes and pathways

In contrast to comparing overall transcriptomic changes between cognitively healthy donors and patients with AD, assessment of transcriptomic changes in individual cell types revealed slightly altered numbers of AD-associated DEGs (Fig 4). The top four most upregulated AD-associated genes found between control and AD libraries (*STC1, MMP1, TFPI2, CBX3*) were shared between all cell types excluding the unknown cells. Only 21 AD-associated DEGs were common to all three cell types (Suppl. table 4). The up-regulated AD-associated genes, *PCSK7, MMP1* and *MMP14*, potentially linked to APP processing, were up-regulated in fibroblast/stromal – and GBC -like cells of OM derived from patients with AD (Fig 3e and Suppl. table 4). In addition, up-regulation of *MMP1* and *MMP14* were detected in myofibroblast-like cells of AD patient OM (Suppl. table 4). Collectively, these three genes are also associated with cytokine signaling in immune system and ECM organization. In AD, fibroblast/stromal – and GBC-like cells are the most transcriptionally active with 80 and 81 up-regulated genes (Fig 4a, b). Figure 4c shows the most up- and down-regulated differentially expressed genes in each cell type. For this the GBC - like cells in different cell cycle phases were considered as one cell type. Interestingly, fibroblast/stromal -like cells exhibited the most unique, AD-associated DEGs (46 genes) (Fig 4b, d & suppl. table 4). Unique for AD GBC -like cells was for example the up-regulation of mitochondrial genes *MT-ND3* and *MT-ND4L*, and down-regulation of *TOMM5*. Highly down-regulated genes selectively in fibroblast/stromal -and myofibroblast -like cells included genes encoding for clusterin (*CLU*) and APP homolog (*APLP2*), and several ribosomal genes, respectively. Using the 147 DEGs significantly up-regulated and down-regulated in AD shown in figure 3e, we then performed pathway analysis with Reactome database and GSVA and identified a total of 73 significantly enriched pathways (Suppl. table 5). Enriched pathways were associated with RNA and protein metabolism, inflammatory processes, signal transduction, cell cycle, developmental biology and the neuronal system (Fig 4d, suppl. fig. 4). Interestingly, our analysis revealed multiple pathways differentially expressed between the AD and control OM cells. For example, pathways related to axon guidance, seleno-amino acid metabolism, selenocysteine synthesis, and ribosomal RNA metabolism were reduced in all AD OM cell types compared to the OM cells derived from cognitively healthy controls.

**Figure 4.**
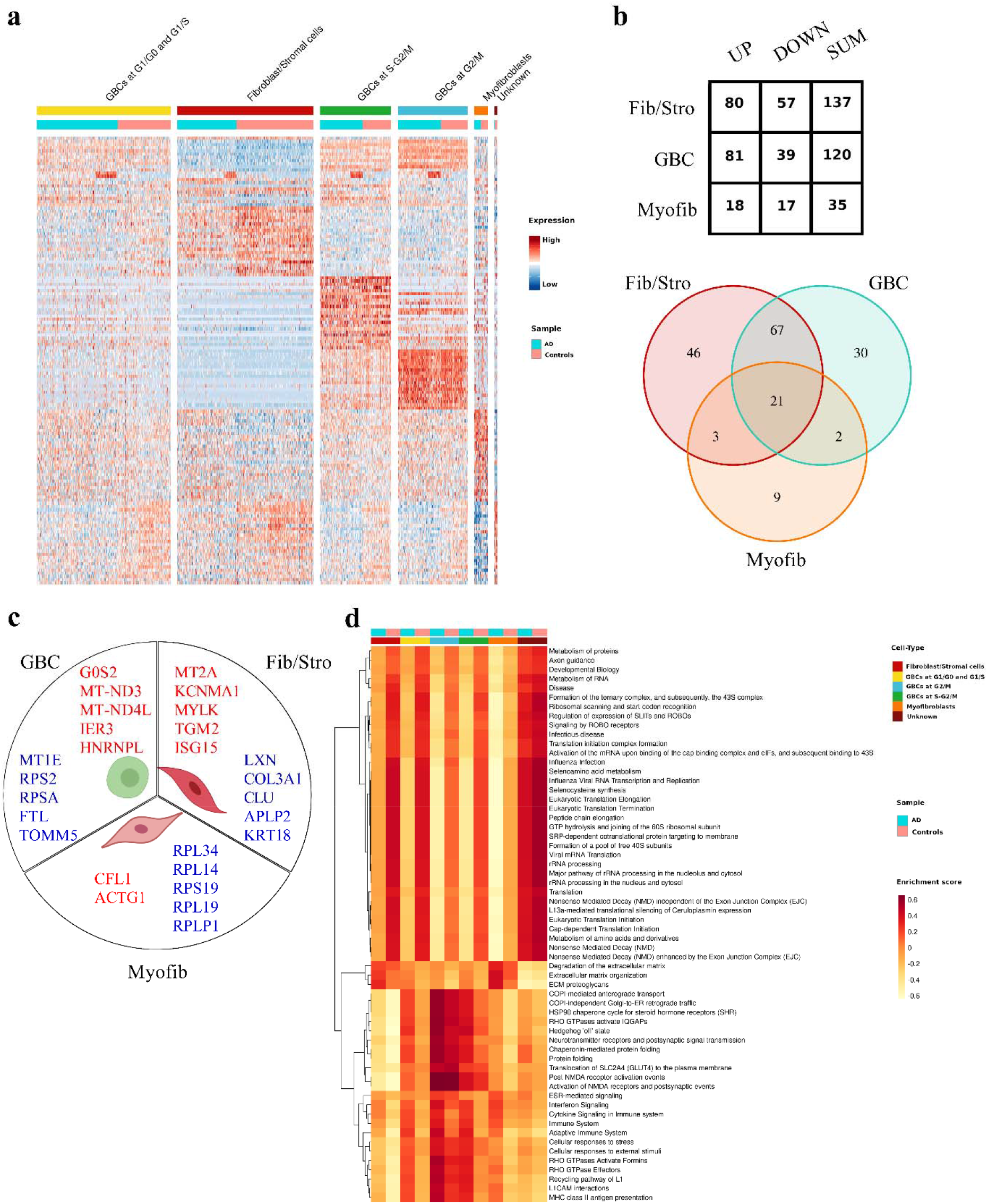
Cell-type-specific pathways altered in AD. **a)** Heatmap depicting differential gene expression at the single-cell level for the most differentially expressed genes for each annotated cell type separated between donors with AD and cognitively healthy individuals. Each expression of the values is scaled and centered in 0. The range of expression values is from −4 to 4. **b)** Numbers of differentially expressed AD-related genes by cell types common for controls and AD shown as table and Venn-diagram. **c)** The top five up-regulated (in red) and down-regulated (in blue) AD-related genes differentially expressed between control and AD OM by cell types. **d)** Heatmap of the subset of differentially expressed pathways for cell types common between AD and controls shows GSVA enrichment score of the pathways obtained for the DEGs with adjusted p-value <0.05. The enrichment score range is from −0.6 to 0.6. GBC, globose basal cell -like cells. Fib/Stro, fibroblast/ stromal -like cells. Myofib, myofibroblast -like cells.

Pathways enriched in GBC -like cells included Hedgehog signaling and Hedgehog “off” state, both of which were more enriched in AD compared to control (for both pathways *TUBA1C*, *TUBA1B*, *PSMD14*, *PSMD2*, *FLNB*, *TUBB4B*). Furthermore, GBC-like cells were enriched for pathways related to neurotransmitter receptors and their postsynaptic effects. Pathways enriched in fibroblast/stromal -and myofibroblast-like cells included ECM degradation (*MMP14*, *MMP1*, *ELN*, *HTRA1*, *COL6A3*, *DCN*, *CTSB*), organization (*MMP14*, *LUM*, *MMP1*, *ELN*, *SERPINE1*, *HTRA1*, *BGN*, *COL6A3*, *THBS1*, *DCN*, *CTSB*), and ECM proteoglycans (*LUM*, *SERPINE1*, *BGN*, *COL6A3*, *DCN*) (Fig 4d).

## DISCUSSION

Olfactory impairment is an early symptom of AD, yet to date, the AD OM remains poorly studied. Although previous reports demonstrated by histological assessment the presence of pathology typical for the disease, cell-type-specific changes in AD OM remain under-explored. Here, we report the OM cell culture as a highly physiologically relevant model to study AD *in vitro*. Our results demonstrate the secretion of toxic Aβ by AD OM cells. Single-cell transcriptomic signatures of these cells revealed 147 differentially expressed AD-associated genes that are primarily involved in pathways related to RNA and protein metabolism, inflammatory processes and signal transduction.

The OM is composed of three primary components, the epithelium, basement membrane, and lamina propria. *In vivo*, the healthy human OM consists of several cell types, including for example olfactory sensory neurons, basal cells (stem cells), Bowman’s gland cells, vascular smooth muscle cells, sustentacular cells and immune cells, as recently described ^16^. Our findings reveal that cultured OM cells can produce and secrete Aβ_1-40_ and Aβ_1-42_, with AD patient cells secreting significantly more Aβ_1-42_ than cognitively healthy control cells. Prior literature demonstrates that the nasal mucosa or nasal secretions of AD patients also exhibit increased levels of Aβ ^7,9–11,20^. Furthermore, the amount of Aβ_1-42_ in the nasal area of transgenic Tg2576 mice modelling AD positively correlates with Aβ_1-42_ deposition in their brain ^21^. The fact that AD OM cells secrete increased levels of toxic Aβ demonstrates that this *in vitro* model shows the expected disease-specific pathological alterations.

Beyond the OM, olfactory sensory cues are processed by the olfactory bulb, olfactory tract, piriform cortex, amygdala, entorhinal cortex, and hippocampus ^1^. ScRNA-seq results by Grubman *et al*. on the AD entorhinal cortex and our findings have the following seven common DEGs between the control and AD groups ^22^: *SERPINE1*, *IFI27*, *MT-ND2*, *MAP1B*, *BCYRN1*, *ANGPTL4* and *HES1*. The presence of the common DEGs highlights the importance of these genes in AD independently of the tissue type assessed and suggests that the entorhinal cortex and OM, both vulnerable to early AD pathogenesis, display disease-specific alterations that may have potential for targeting by therapeutic approaches.

To deduce why Aβ_1-42_ secretion is increased in AD OM cells, we found the expression of both genes *PCSK7* and *MMP14* to be up-regulated in fibroblast/stromal – and GBC -like cells (logFC 0.41 and 0.35 for *PCSK7* and logFC 0.40 for MMP14 in both cell types, respectively). Our pathway analysis of the 90 up-regulated genes in the AD OM cells revealed PCSK7, and MMP14 to be involved Alzheimer’s disease-presenilin pathway (PANTHER pathway P00004). Both *PCSK7* and *MMP14* are linked to APP processing ^23–25^, although their role in APP processing in human OM has not been studied. PCSK7 is activated by metallothionein 3 (*MT3*), which in turn activates APP cleaving α-secretase and metalloprotease, ADAM10 ^25^. Furthermore, as a possibly unique feature of APP processing in the human OM, we observed reduced expression of genes encoding for secreted frizzled related proteins in AD OM fibroblast/stromal – and GBC - like cells compared to those of the control OM (*SFRP1* logFC −0.64 and −0.83, *SFRP2* logFC −1.54 and −0.72, respectively). In brains affected by AD, SFRP1 is known to be elevated and promote pro-amyloidogenic APP processing by negatively regulating the α-secretase ADAM10 activity ^26^. Further in-depth experiments are needed to study the role of SFRPs on APP processing in the human OM.

Our results indicated that the gene expression level of *MT2A* (metallothionein 2A) was significantly and selectively up-regulated (logFC 0.56) in AD OM derived fibroblast/stromal – like cells while no changes in *MT3* expression were observed. Previous studies have also demonstrated an increase in MT2A expression in skin fibroblasts of AD patients ^27^, supporting our observation. Interestingly, the gene encoding for metallothionein 1E (*MT1E)* was significantly down-regulated solely in GBC -like cells (logFC −0.33) derived from AD OM. The expression of metallothionein proteins are suggested to become elevated in response to increased amounts of metals or reactive oxygen species (ROS) in the cell (for review see ^28^). On the other hand, when MT2A is overexpressed in HEK cells, the cells display reduced capacity for oxidative phosphorylation, thereby affecting their mitochondrial function ^29^. These findings suggest that alterations in mitochondrial function may occur in cultured AD OM cells in a similar manner as observed in other cells in AD (reviewed in ^30,31^). This idea is supported by the finding that GBC -like cells derived from the AD OM showed selective up-regulation of complex I subunits *MT-ND3* and *MT-ND4L* simultaneously with down-regulation of the gene encoding for the outer mitochondrial membrane protein TOMM5. However, genes of the complex I are reported to be down-regulated in AD brains ^32,33^. Detailed studies into the effects of the heavy metal binding proteins on APP processing and mitochondrial function in OM would form an interesting future direction of research.

Although hyperphosphorylated tau was detected in neurites in AD OM histological sections ^10^, we did not observe a change in levels of total or phosphorylated tau between AD and control OM cells. This lack of change is not surprising given that mature neurons were not detected in our OM cell cultures, where tau is expressed endogenously ^34^.

Multiple genome-wide association studies have found pathways related to metabolism and immune system to be altered in AD ^35–37^, thus supporting our findings of altered DEGs related to pathways including RNA and protein metabolism, inflammatory processes, signal transduction, and developmental biology. We observed that immune system pathways were significantly enriched in the AD OM cells, which also aligns with earlier published studies ^38,39^. Clusterin (encoded by *CLU*), also known as apolipoprotein J, is strongly linked to AD and up-regulated in AD patient brains as a protective response (for review see ^40^). Interestingly, in our study, *CLU* was selectively down-regulated in AD OM derived fibroblast/stromal -like cells. Prior studies have shown that in lung fibroblasts *CLU* expression is reduced by transforming growth factor β 1 (TGFβ_1_) ^41^, which is in contrast to neurons and astrocytes, where TGFβ induces clusterin expression (for review see ^41^). It is therefore possible that clusterin alterations in AD are cell-type specific and related to TGFβ levels. Furthermore, it is known that the fibroblast proliferation promoting effects of serum are diminished in cells expressing low levels of *CLU* ^41^. This, together with our results of reduced numbers of fibroblast/stromal cells in AD OM cultures grown in serum-containing media, may suggest the involvement of clusterin in proliferation of OM-derived fibroblast/stromal cells *in vitro*.

Although our CBA results did not reflect altered cytokine secretion in AD OM cells, the pathway analysis revealed that AD-associated DEGs enrich for Cytokine Signaling in Immune System, Interferon Signaling, Adaptive Immune System, and Signaling by Interleukins. As resilient chronic inflammation is considered as a renowned hallmark of AD ^6^, longer culture periods or exposure to inflammatory stimuli could reveal differential cytokine profiles between the AD and control OM cultures, as previously demonstrated in healthy OM cells exposed to air pollutants ^42,43^.

In the cell cultures we identified fibroblast/ stromal -like cells, myofibroblast-like cells, and GBC -like cells as cell types dominantly present in OM of cognitively healthy controls and AD patients. Although some cell types remained unidentified, it is likely that the cultures do not contain all the cell types present in the OM *in vivo*. This is to be expected given that the more robust cell types such as fibroblast/ stromal -like cells tend to survive the culturing conditions better and dominate the more sensitive and low abundance cells such as sensory neurons. Furthermore, the culture conditions, including culture media composition, is likely to better promote the growth of some cell types while lacking support for others. To mention one, Newman *et al.* have shown that neurons do not survive the primary cultures of OM cells derived from adult mice, but they can be induced by a right cocktail of growth factors ^44^. While our data demonstrated that donor to donor variation was minimal with regards to the proportions of cell types present, the proportions of the GBC -like cells were slightly increased in cultures derived from patients with AD when compared to cognitively healthy individuals. On the other hand, the control OM cultures contained more fibroblast/ stromal cell -like cells by proportion compared to AD OM cultures. However, the AD GBC -like cells enriched more in pathways related to hedgehog signaling compared to other cell types or control GBC-like cells, potentially resulting from enhanced proliferation and negative regulation of Wnt signaling, mediated by up regulation of *SFRP1* (secreted frizzled related protein 1) observed in AD OM cells (for review see ^45^).

To open new avenues for therapy and to overcome the failure of primarily neuron-targeting treatments for AD, current research focus has diverted towards non-neuronal cells. Given that the OM cells are easily accessible from living donors, this enables repeated assessment of the disease-associated features during disease progression. In addition, these cells could be highly useful for diagnostic purposes when harvested from individuals at early disease stages. The OM cells also reflect the heterogeneity of genetic risk, such as aged patients with LOAD, when cells from multiple patients are included in the experiments. While several studies have produced evidence that the widely used induced pluripotent stem cells (iPSCs) lines can acquire a variety of genetic and epigenetic aberrations during the reprogramming process ^46^, the OM cells retain epigenetic alterations obtained throughout life ^47^. Furthermore, 3D culture may promote even stronger AD-related disease phenotypes in the OM cells, as has been demonstrated in iPSC-derived co-cultures (for review see ^46^). The results shown here demonstrate that the OM cells present high research potential in complex age-related diseases, such as LOAD, and present evidence for their ability to reveal specific molecular mechanisms and cellular processes that could reveal new targets for diagnosis and therapeutic intervention.

## METHODS

### Ethical considerations

The study was performed with the approval of the Research Ethics Committee of the Northern Savo Hospital District (permit number 536/2017). The oral and written information concerning the study was provided by the study investigator or study nurse. The written informed consent was collected from all the voluntary study participants and proxy consent from the family members /legally acceptable representatives of persons with mild AD dementia.

### Patients and OM cell cultures

A total of 12 voluntary study patients with AD type mild (CDR 1) dementia were recruited via Brain Research Unit, Department of Neurology, University of Eastern Finland. AD diagnostic examinations had been carried out at Brain Research Unit or at the Department of Neurology, Kuopio University Hospital prior to study recruitment. All the patients with AD dementia fulfilled the NIA-AA clinical criteria of progressive AD and MRI imaging or FDG-PET study had showed degenerative process or, in CSF biomarker examination was found biomarker (beta-amyloid, tau and phos-tau) changes typical to AD ^48^. Eleven cognitively healthy control subjects were recruited via Department of Otorhinolaryngology, Kuopio University Hospital, Finland from patients undergoing a dacryocystorhinostomy (DCR) surgery, or from the already existing registries of the Brain Research Unit of University of Eastern Finland. Cognition of all the study patients were evaluated utilizing the Consortium to Establish a Registry for Alzheimer’s Disease (CERAD) neuropsychological battery ^49,50^. The age of the patients with AD and cognitively healthy control subjects age average 68.3 years and 70.6 years, respectively. For AD group 50% of the study subjects were males and 50% females, whereas for control subjects 27.3% were males and 72.7% females. A venous blood sample was taken from all study participants for use in *APOE*-genotyping. Based on genotyping, 66.7% of the patients with AD and 36.4% of cognitively healthy control subjects had at least one *APOE* ε4 allele. Patients and cognitively healthy study participants were tested for their sense of smell for 12 odors (Sniffin’ Sticks, Heinrich Burghart GmbH, Wedel, Germany) and classified as normal, hyposmic, or anosmic ^51^.

A piece of the OM was collected as a biopsy from the nasal septum, close to the roof of the nasal cavity as previously described ^2,52^ The biopsy was kept on ice in DMEM/F12 (#11320033) based growth medium containing 10% heat -inactivated fetal bovine serum (FBS) (#10500064) and 1x Penicillin-Streptomycin solution (#15140122) until processed in the laboratory (All reagents Thermo Fisher Scientific, Waltham, MA, USA). Primary OM cell cultures were established according to the published protocol ^53,54^ with small modifications. In short, any remaining of blood or cartilage were removed under a dissection microscope and the flattened tissue piece was rinsed several times with cold Hank’s Balanced Salt Solution. To separate olfactory epithelium and underlying lamina propria, the tissue was enzymatically digested for 45 minutes with dispase II as 2.4 U/ml (Roche, Basel, Switzerland) followed by further digesting the lamina propria with DMEM/F12 media containing 0.25 mg/ml collagenase H (Sigma-Aldrich, St. Louis, MO, USA) for up to 10 minutes. Finally, the digested olfactory epithelium and lamina propria were combined and seeded on poly-D-lysine (Sigma-Aldrich) coated 6-well in order to let the cells migrate out of the tissue pieces and proliferate in growth medium at 37°C, 5% CO_2_. Half of the culture media was changed every 2-3 days for in total of 8 to 19 days before passaging the cultures and freezing the primary cell lines in liquid nitrogen for later use in solution containing 90 % heat inactivated FBS and 10 % dimethyl sulfoxide. Cells in primary passages of 2-3 were used for scRNA-seq and primary passages 4-6 for biochemical analyses.

### *APOE*-genotyping

*APOE*-genotyping of the study subjects was performed as described previously ^53,54^. Briefly, the genomic DNA was isolated from venous blood samples with QIAamp DNA blood mini extraction kit (QIAGEN) and the *APOE* alleles were determined by polymerase chain reaction for two single-nucleotide polymorphisms (rs429358 and rs7412) with Taqman SNP Genotyping Assays (Applied Biosystems, Foster City, CA, USA). The polymerase chain reaction and following allelic discrimination were performed on QuantStudio 5 Real-Time PCR System platform (Applied Biosystems).

### Enzyme-linked immunosorbent assays

The culture medium was collected and cells were lysed in RIPA buffer supplemented with 1x cOMPLETE protease inhibitor cocktail (Roche, Basel, Switzerland). Cell lysates and medium for phosphorylated tau and total tau Enzyme-linked immunosorbent assays (ELISA) were collected in RIPA buffer supplemented with 1x cOmplete protease inhibitor cocktail as well as phosphatase inhibitor cocktail 2 at 1:100 dilution (#P5726, Sigma-Aldrich, MO, USA). Samples were stored at −70°C until analysis. 50 μl of each medium, cell lysate or plasma sample was analyzed in singlets for cell lysates and media samples, and in duplicates for plasma samples using ELISA kits (all from Invitrogen, CA, USA) for human Amyloid beta 40 (#KHB3481), Amyloid beta 42 (#KHB3544), tau (phospho, pT181) (#KHO0631) and tau (total) (#KHB0041), according to the manufacturer’s instructions. Total protein amounts of the OM cell lysates were quantified with BCA assay (Pierce™ BCA Protein Assay Kit, Thermo Scientific) according to the manufacturer’s instructions. The results were calculated as pg/mg protein ± SD for the OM cells and for plasma samples as pg/ml plasma ± SD.

### Cytometric Bead Array for Cytokine Secretion

The cell culture medium was collected, and cells were lysed in RIPA buffer supplemented with 1x cOmplete protease inhibitor cocktail (Roche, Basel, Switzerland). 20 μl of each sample was analysed in duplicates using the BD™ CBA Human Soluble Protein Master Kit (#556264; BD Biosciences, CA, USA) alongside the BD™ CBA Human Flex Sets for interleukin 1ß (IL-1ß; #558279), interleukin 6 (IL-6; #558276), interleukin 8 (IL-8; #558277), monocyte chemoattractant protein 1 (MCP-1; #558287), tumour necrosis factor (TNF, #560112). Data were acquired using CytoFLEX S (Beckman Coulter Life Sciences, IN, USA) and analysed with FCAP Array™ v2.0.2 software (Soft flow Inc, MN, USA). Protein concentrations were quantified with BCA kit from the cell lysates as described for the ELISA assays and results were calculated as pg/μg total protein ± SD.

### Cell hashing and single cell RNA sequencing

Cells from three cognitively healthy control lines (average age 71.7 years) and two AD patients (average age 72 years), all females with *APOE33* genotype, were harvested from culture with TrypLE Express (Gibco, CA, USA) and resuspended in PBS containing 2% BSA. TotalSeq™ -A anti-human Hashtag Antibodies (BioLegend, San Diego, CA, USA) were used according to the producer’s instructions to stain the cells of individual donors before pooling the cells as one pool for control subjects and one pool for AD patients. Equal numbers of cells from each donor were pooled together. Next, the cell pools were filtered through a 30 μm strainer and the cells were resuspended in PBS containing 0.04% BSA. Viability of the single-cell suspension pools was > 90% based on Trypan blue staining.

The sample pools were run on individual lanes of a Chromium Chip B with the Chromium Single Cell 3′ Library & Gel Bead Kit v3 kit (10x Genomics, CA, USA) according to the manufacturer’s protocol with a targeted cell recovery of 10,000 cells per lane. In the cDNA amplification step, the cell hashing protocol from New York Genome Center Technology Innovation Lab (version 2019-02-13) was followed (https://genomebiology.biomedcentral.com/articles/10.1186/s13059-018-1603-1). The hashtag oligonucleotide primer (HTO primer 5’GTGACTGGAGTTCAGACGTGTGCTC’3) was added to the cDNA amplification mix. During cDNA cleanup, the supernatant contained the HTO-derived cDNAs (<180bp) and the pellet the mRNA-derived cDNAs (>300bp). The mRNA-derived cDNAs were processed according to the protocol provided by 10x Genomics and the HTO-derived cDNAs fraction was processed according to the cell hashing protocol referenced above. The 3’ gene expression libraries and HTO sequencing libraries were pooled and sequenced at an approximate depth of 50,000 reads per cell for the 3’ gene expression libraries and 5,000 reads for HTO libraries using the NovaSeq S1 (Illumina, San Diego, CA, USA) flow cells.

### Quality control and downstream analysis

Cell Ranger v.3.0.2 was used to analyze the raw base call files.□ FASTQ files and raw gene-barcode matrices were generated using the command ‘mkfastq’ and the ‘count’ command, respectively, and aligned human genome GRCh37 (hg19).□ The samples were integrated in R v.4.0.3□and the two generated Seurat objects, one related to Alzheimer’s disease samples and the other to control samples, were analyzed using the Seurat package v.3.2 ^53,54^.□ Working separately on the Controls and Alzheimer’s disease samples dataMatrix, hashtag oligos assays were normalized with NormalizeData() function applying the centered log-ratio (CLR) transformation. Seurat’s HTODemux was ran with default parameters on the hashtag-count matrices to demultiplex interested cell barcodes. HTO count distribution was checked and cells classified into singlet/doublet/negative. With HTODemux single cells were assigned back to their samples of origin. After the demultiplexing step, the cells filtering was performed separately for the two libraries removing low quality cells and keeping those with nFeature > 750 and nFeature < 7000, nCount_RNA < 40000 and mitochondrial content < 12%. Finally, doublets were removed using the subset() function. Control and Alzheimer’s disease samples were integrated using Seurat standard integration pipeline. The function SplitObject() was used to separate, inside the two libraries, each sample which was, except for negative cells omitted in the integration process, individually normalized using the NormalizeData() function and the standard ‘LogNormalize’ method. From each of the five samples the most 5000 variable features were extracted using the FindVariableFeatures() function applying the ‘vst’ method. Thereafter, all the samples were integrated, at first by extracting the anchors between samples using 5000 anchor features and 30 dimensions as parameters in the FindIntegrationAnchors() function, then, with the IntegrateData() function all the five samples were integrated using the previously extracted anchors and as before the first 30 dimensions. The resulting Seurat object was then scaled with ScaleData() function and PCA was applied using RunPCA() function. With the FindNeighbors() function, using the first 30 dimensions, the Shared Nearest Neighbor Graph was constructed and cells clustered using the FindClusters() function. Different clustering resolutions were tested.

The cluster resolution value 0.2 was the best to represent cell-type annotation. RunUMAP() function was used to apply UMAP dimensional reduction and obtain a spatial visualization of the cells. For each clusters the top 30 over-expressed genes were identified with FindAllMarkers() function using the Wilcox rank sum test and setting the min.pct and logfc.threshold parameters to 0.3. The significant genes were inputted, for the cell-type annotation, into EnrichR ^55,56^ and HumanBase ^57^ tools, using Gene Ontology and cellular pathway information to verify the output results. In order to obtain more markers for the annotation than those found initially, cluster 0 was also analyzed separately with a resolution of 0.2. To be more confident of the final cell-type annotation, we also considered the effects of cell cycle heterogeneity in our data calculating the cell cycle phase score for each cell using CellCycleScoring() function provided by Seurat package.

### Differential expression and enrichment

The differential expression was performed between Alzheimer’s disease and Controls samples and also between cells in each cell-type subpopulation. FindMarkers() function was used. In order to mitigate batch effect, MAST test was applied using the samples of origin as latent variable parameter. To understand in which biological processes the differential expressed genes are involved, enrichment analysis was performed using a non-parametric unsupervised method called Gene Set Variation Analysis (GSVA, v.1.38.0) ^58^ and the open-source pathway database Reactome ^59^.

### Statistical methods and graphical illustrations

GraphPad Prism 8.1.0 (GraphPad Software Inc. San Diego, CA, USA) software was used for statistical analysis of the data. Mean values in ELISA and CBA analyses, between control and AD, was compared using unpaired two-tailed t-test with Welch’s correction or with two-way ANOVA. Error bars in the figure legends represent standard deviation (SD). Statistical significance was assumed for *p*-values < 0.05. The graphical illustrations were created with BioRender.com and open source vector graphics editor Inkscape 0.91.

## Acknowledgements

The authors would like to thank Ratneswary Sutharsan for assistance in establishing the OM culture conditions, Virve Kärkkäinen for help with ethical aspects, and Mirka Tikkanen for technical assistance. We gratefully acknowledge the support of The Academy of Finland, The Juselius Foundation, The Finnish Cultural Foundation, The Inkeri and Mauri Vänskä Foundation, The Yrjö Jahnsson Foundation, The Saastamoinen Foundation and The Finnish Brain Foundation. This project has received funding from the European Union’s Horizon 2020 research and innovation programme under grant agreement No 814978. Sequencing was performed by the Sequencing unit of Institute for Molecular Medicine Finland FIMM Technology Centre, University of Helsinki. Sequencing unit it supported by Biocenter Finland.

## Contributions

These authors contributed equally: Riikka Lampinen, Mohammad Feroze Fazaludeen K.M.K., A.R.W., A.M.K., T.M., A.M-S., and H.L., conceived the project. K.M.K., M.U.K., T.Ö., R.L., and S.C. designed the scRNA-sequencing experiment. R.L. and M.F.F. performed scRNA-seq library generation. R.G., S.A., M.F.F., T. Ö., and R.L. performed data analysis with assistance from S.C., L.S., and E.K. S.C., R.L., M.C-H., and F.F.A. performed the cell functional assays. E.P., S.H., T.S., J-M.L., and A.M.K. performed clinical assessments of the study subjects. R.L., S.C., M.F.F., and K.M.K. wrote the manuscript. A.M-S. and A.M.K. revised the manuscript.

## Corresponding author

Correspondence and requests for materials should be addressed to Katja M Kanninen.

## Supplementary Information

Supplementary Information is available for this paper.

## REFERENCES

1. Marin, C. et al. Olfactory Dysfunction in Neurodegenerative Diseases. Current Allergy and Asthma Reports 18, 42 (2018).

2. Féron, F., Perry, C., McGrath, J. J. & Mackay-Sim, A. New techniques for biopsy and culture of human olfactory epithelial neurons. Archives of Otolaryngology - Head and Neck Surgery 124, 861–866 (1998).

3. Jung, H. J., Shin, I.-S. & Lee, J.-E. Olfactory function in mild cognitive impairment and Alzheimer’s disease: A meta-analysis. The Laryngoscope 129, 362–369 (2019).

4. Sohrabi, H. R. et al. Olfactory dysfunction is associated with subjective memory complaints in community-dwelling elderly individuals. Journal of Alzheimer’s Disease 17, 135–142 (2009).

5. Sohrabi, H. R. et al. Olfactory discrimination predicts cognitive decline among community-dwelling older adults. Translational Psychiatry 2, (2012).

6. Lane, C. A., Hardy, J. & Schott, J. M. Alzheimer’s disease. European Journal of Neurology 25, 59–70 (2018).

7. Kim, Y. H. et al. Amyloid beta in nasal secretions may be a potential biomarker of Alzheimer’s disease. Scientific Reports 9, 4966 (2019).

8. Liu, Z. et al. Development of a High-Sensitivity Method for the Measurement of Human Nasal Aβ 42, Tau, and Phosphorylated Tau. Journal of Alzheimer’s Disease 62, 737–744 (2018).

9. Yoo, S. J. et al. Longitudinal profiling of oligomeric Aβ in human nasal discharge reflecting cognitive decline in probable Alzheimer’s disease. Scientific Reports 10, 11234 (2020).

10. Arnold, S. E. et al. Olfactory epithelium amyloid-β and paired helical filament-tau pathology in Alzheimer disease. Annals of Neurology 67, 462–469 (2010).

11. Ayala-Grosso, C. A. et al. Amyloid-Aβ peptide in olfactory mucosa and mesenchymal stromal cells of mild cognitive impairment and Alzheimer’s disease patients. Brain Pathology 25, 136–145 (2015).

12. Talamo, B. R. et al. Pathological changes in olfactory neurons in patients with Alzheimer’s disease. Nature 337, 736–739 (1989).

13. Lee, J. H., Goedert, M., Hill, W. D., Lee, V. M. Y. & Trojanowski, J. Q. Tau Proteins Are Abnormally Expressed in Olfactory Epithelium of Alzheimer Patients and Developmentally Regulated in Human Fetal Spinal Cord. Experimental Neurology 121, 93–105 (1993).

14. Ghanbari, H. A. et al. Oxidative damage in cultured human olfactory neurons from Alzheimer’s disease patients. Aging Cell 3, 41–44 (2004).

15. Wolozin, B., Lesch, P., Lebovics, R. & Sunderland, T. Olfactory neuroblasts from Alzheimer donors: Studies on APP processing and cell regulation. Biological Psychiatry 34, 824–838 (1993).

16. Durante, M. A. et al. Single-cell analysis of olfactory neurogenesis and differentiation in adult humans. Nature Neuroscience 23, 323–326 (2020).

17. Holbrook, E. H., Rebeiz, L. & Schwob, J. E. Office-based olfactory mucosa biopsies. International Forum of Allergy and Rhinology 6, 646–653 (2016).

18. Mi, H., Muruganujan, A., Ebert, D., Huang, X. & Thomas, P. D. PANTHER version 14: More genomes, a new PANTHER GO-slim and improvements in enrichment analysis tools. Nucleic Acids Research 47, D419–D426 (2019).

19. Mi, H. & Thomas, P. PANTHER pathway: an ontology-based pathway database coupled with data analysis tools. Methods in molecular biology (Clifton, N.J.) 563, 123–140 (2009).

20. Liu, Z. et al. Development of a High-Sensitivity Method for the Measurement of Human Nasal Aβ 42, Tau, and Phosphorylated Tau. Journal of Alzheimer’s Disease 62, 737–744 (2018).

21. Kameshima, N., Nanjou, T., Fukuhara, T., Yanagisawa, D. & Tooyama, I. Correlation of Aβ deposition in the nasal cavity with the formation of senile plaques in the brain of a transgenic mouse model of Alzheimer’s disease. Neuroscience Letters 513, 166–169 (2012).

22. Grubman, A. et al. A single-cell atlas of entorhinal cortex from individuals with Alzheimer’s disease reveals cell-type-specific gene expression regulation. Nature Neuroscience 22, 2087–2097 (2019).

23. Lopez-Perez, E., Seidah, N. G. & Checler, F. Proprotein Convertase Activity Contributes to the Processing of the Alzheimer’s β-Amyloid Precursor Protein in Human Cells: Evidence for a Role of the Prohormone Convertase PC7 in the Constitutive -Secretase Pathway. Journal of Neurochemistry 73, 2056–2072 (2002).

24. Py, N. A. et al. Differential spatio-temporal regulation of MMPs in the 5xFAD mouse model of Alzheimerâ€^™^s disease: evidence for a pro-amyloidogenic role of MT1-MMP. Frontiers in Aging Neuroscience 6, 247 (2014).

25. Park, B. H., Kim, H. G., Jin, S. W., Song, S. G. & Jeong, H. G. Metallothionein-III increases ADAM10 activity in association with furin, PC7, and PKCα during non-amyloidogenic processing. FEBS Letters 588, 2294–2300 (2014).

26. Esteve, P. et al. Elevated levels of Secreted-Frizzled-Related-Protein 1 contribute to Alzheimer’s disease pathogenesis. Nature Neuroscience 22, 1258–1268 (2019).

27. Lanni, C. et al. Homeodomain Interacting Protein Kinase 2: A Target for Alzheimer’s Beta Amyloid Leading to Misfolded p53 and Inappropriate Cell Survival. PLoS ONE 5, e10171 (2010).

28. Lindeque, J. Z., Levanets, O., Louw, R. & van der Westhuizen, F. H. The Involvement of Metallothioneins in Mitochondrial Function and Disease. Current Protein & Peptide Science 11, 292–309 (2010).

29. Bragina, O. et al. Metallothionein 2A affects the cell respiration by suppressing the expression of mitochondrial protein cytochrome c oxidase subunit II. Journal of Bioenergetics and Biomembranes 47, 209–216 (2015).

30. Chakravorty, A., Jetto, C. T. & Manjithaya, R. Dysfunctional Mitochondria and Mitophagy as Drivers of Alzheimer’s Disease Pathogenesis. Frontiers in Aging Neuroscience 11, 311 (2019).

31. Wang, W., Zhao, F., Ma, X., Perry, G. & Zhu, X. Mitochondria dysfunction in the pathogenesis of Alzheimer’s disease: Recent advances. Molecular Neurodegeneration 15, 30 (2020).

32. Manczak, M., Park, B. S., Jung, Y. & Reddy, P. H. Differential Expression of Oxidative Phosphorylation Genes in Patients with Alzheimer’s Disease: Implications for Early Mitochondrial Dysfunction and Oxidative Damage. NeuroMolecular Medicine 5, 147–162 (2004).

33. Fukuyama, R., Hatanpää, K., Rapoport, S. I. & Chandrasekaran, K. Gene expression of ND4, a subunit of complex I of oxidative phosphorylation in mitochondria, is decreased in temporal cortex of brains of Alzheimer’s disease patients. Brain Research 713, 290–293 (1996).

34. Kovacs, G. G. Astroglia and Tau: New Perspectives. Frontiers in Aging Neuroscience 12, 96 (2020).

35. Lambert, J. C. et al. Implication of the immune system in Alzheimer’s disease: evidence from genome-wide pathway analysis. Journal of Alzheimer’s Disease 20, 1107–1118 (2010).

36. Jones, L. et al. Correction: Genetic Evidence Implicates the Immune System and Cholesterol Metabolism in the Aetiology of Alzheimer’s Disease. PLoS ONE 6, e13950. (2011).

37. Jiang, Q. et al. Alzheimer’s Disease Variants with the Genome-Wide Significance are Significantly Enriched in Immune Pathways and Active in Immune Cells. Molecular Neurobiology 54, 594–600 (2017).

38. Pillai, J. A. et al. Key inflammatory pathway activations in the MCI stage of Alzheimer’s disease. Annals of Clinical and Translational Neurology 6, 1248–1262 (2019).

39. Chen, J. et al. Gene expression analysis reveals the dysregulation of immune and metabolic pathways in Alzheimer’s disease. Oncotarget 7, (2016).

40. Nuutinen, T., Suuronen, T., Kauppinen, A. & Salminen, A. Clusterin: A forgotten player in Alzheimer’s disease. Brain Research Reviews 61, 89–104 (2009).

41. Peix, L. et al. Diverse functions of clusterin promote and protect against the development of pulmonary fibrosis. Scientific Reports 8, 1–15 (2018).

42. Chew, S. et al. Urban air particulate matter induces mitochondrial dysfunction in human olfactory mucosal cells. Particle and Fibre Toxicology 17, 18 (2020).

43. Osmond-McLeod, M. J. et al. Surface coatings of ZnO nanoparticles mitigate differentially a host of transcriptional, protein and signalling responses in primary human olfactory cells. Particle and Fibre Toxicology 10, (2013).

44. Newman, M. P., Féron, F. & Mackay-Sim. Growth factor regulation of neurogenesis in adult olfactory epithelium. Neuroscience 99, 343–50 (2000).

45. Ding, M. & Wang, X. Antagonism between hedgehog and wnt signaling pathways regulates tumorigenicity (Review). Oncology Letters 14, 6327–6333 (2017).

46. Penney, J., Ralvenius, W. T. & Tsai, L. H. Modeling Alzheimer’s disease with iPSC-derived brain cells. Molecular Psychiatry vol. 25 148–167 (2020).

47. Vitale, A. M. et al. DNA methylation in schizophrenia in different patient-derived cell types. NPJ Schizophrenia 3, 6 (2017).

48. Jack, C. R. et al. NIA-AA Research Framework: Toward a biological definition of Alzheimer’s disease. Alzheimer’s and Dementia 14, 535–562 (2018).

49. Morris, J. C. et al. The consortium to establish a registry for alzheimer’s disease (CERAD). Part I. Clinical and neuropsychological assessment of alzheimer’s disease. Neurology 39, 1159–1165 (1989).

50. Mirra, S. S. et al. The consortium to establish a registry for Alzheimer’s disease (CERAD). Part II. Standardization of the neuropathologic assessment of Alzheimer’s disease. Neurology 41, 479–486 (1991).

51. Hummel, T., Konnerth, C. G., Rosenheim, K. & Kobal, G. Screening of olfactory function with a four-minute odor identification test: Reliability, normative data, and investigations in patients with olfactory loss. Annals of Otology, Rhinology and Laryngology 110, 976–981 (2001).

52. Murrell, W. et al. Multipotent stem cells from adult olfactory mucosa. Developmental Dynamics 233, 496–515 (2005).

53. Stuart, T. et al. Comprehensive Integration of Single-Cell Data. Cell 177, 1888–1902.e21 (2019).

54. Butler, A., Hoffman, P., Smibert, P., Papalexi, E. & Satija, R. Integrating single-cell transcriptomic data across different conditions, technologies, and species. Nature Biotechnology 36, 411–420 (2018).

55. Chen, E. Y. et al. Enrichr: interactive and collaborative HTML5 gene list enrichment analysis tool. BMC Bioinformatics 14, 128 (2013).

56. Kuleshov, M. v. et al. Enrichr: a comprehensive gene set enrichment analysis web server 2016 update. Nucleic acids research 44, W90–W97 (2016).

57. Greene, C. S. et al. Understanding multicellular function and disease with human tissue-specific networks. Nature Genetics 47, 569–576 (2015).

58. Hänzelmann, S., Castelo, R. & Guinney, J. GSVA: gene set variation analysis for microarray and RNA-Seq data. BMC Bioinformatics 14, 7 (2013).

59. Jassal, B. et al. The reactome pathway knowledgebase. Nucleic Acids Research 48, D498–D503 (2020).

