## Supplementary figures 1-4 for "Disease specific alterations in the olfactory mucosa of patients with Alzheimer’s disease"

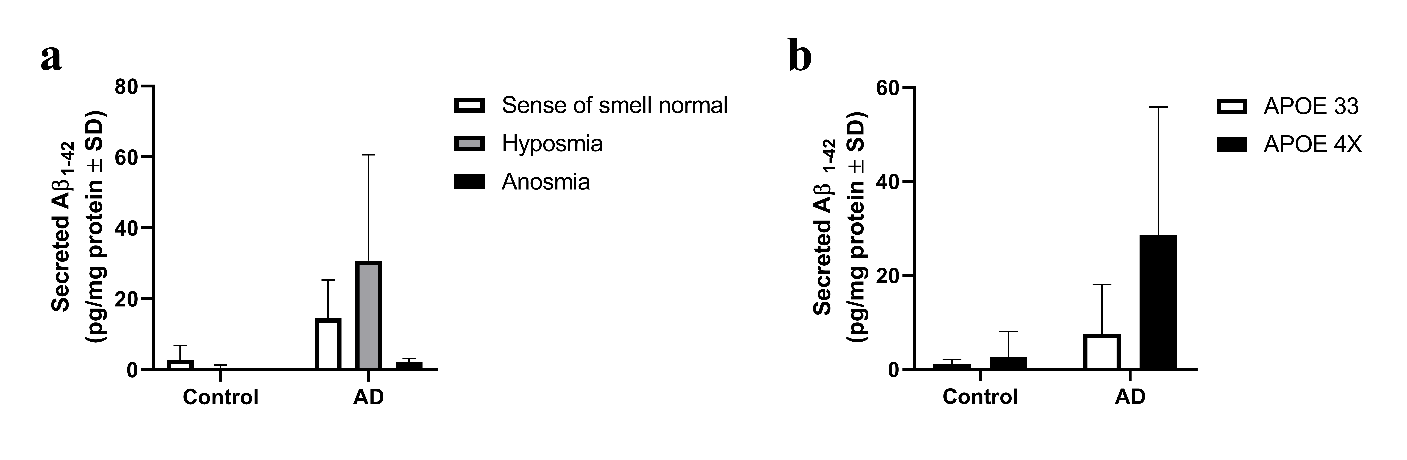

***Supplementary figure 1.*** *OM cells harvested from biopsies were cultured for 7 days prior ELISA assay, assessing levels of Aβ_1-42_ in media collected from OM cells. The results were normalized to the total amount of protein measured from cell lysates and then separated to subgroups based on* ***a)*** *the sense of smell status of the biopsy donor or* ***b)*** *the donor’s APOE genotype. N= 11 donors in total for controls and 10 for AD.* *Quantification of secreted Aβ_1-42_ between control and AD subgroups was performed with two-way ANOVA. A statistically significant difference was detected between the control and AD in the amount of secreted Aβ_1-42_ (p= 0.0338), but not between the APOE genotypes. Data is presented as mean values ± SD.*

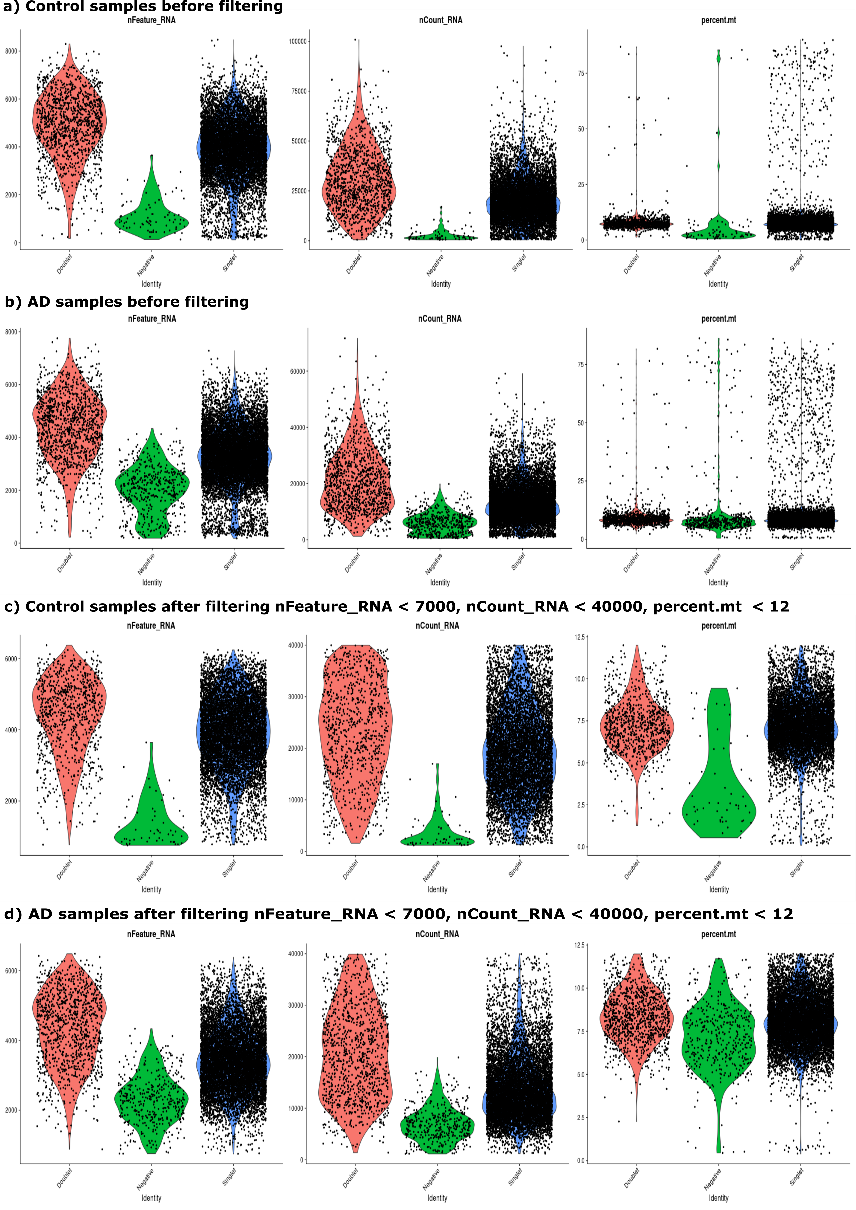

***Supplementary figure 2. Quality control for scRNA-seq data*** *before filtering for a) control library and for b) AD library, and in addition after filtering c) for control library and d) AD library.*

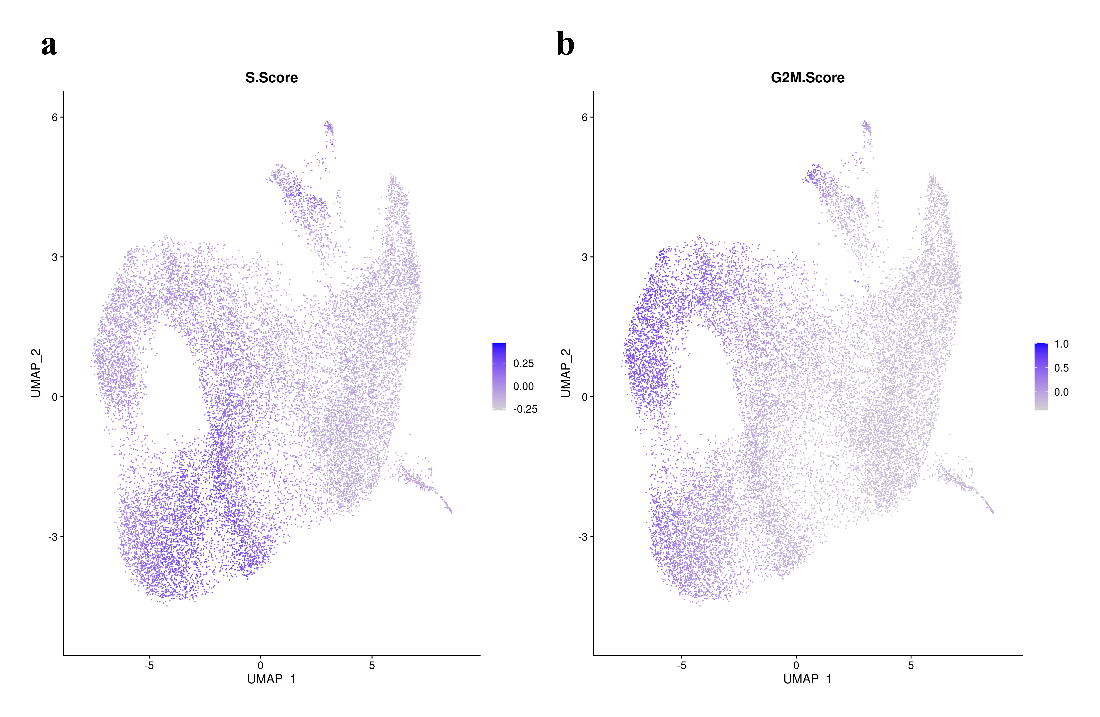

***Supplementary figure 3****.* ***Cell cycle scores based on the expression of G2/M and S phase markers in each cell.*** *The scores have been estimate using the Seurat package.*

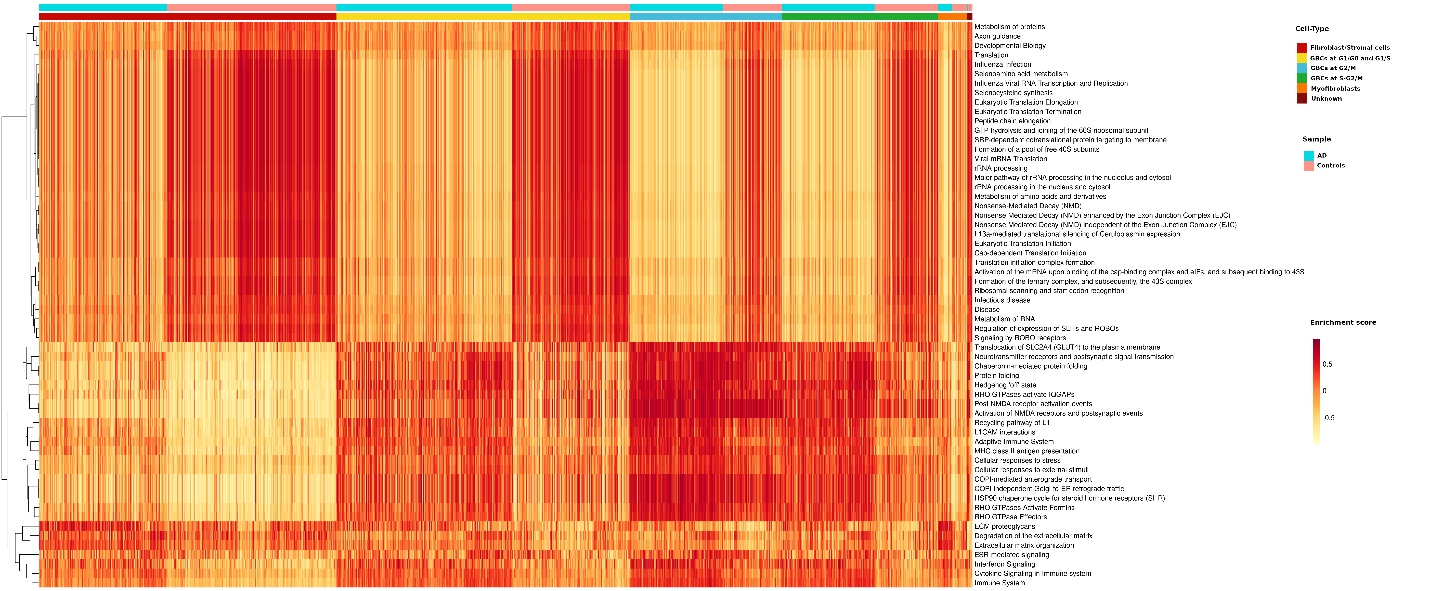

***Supplementary figure 4.*** ***Heatmap of the subset of differentially expressed pathways for cell-types common between AD and controls at single-cell level.*** *The figure shows the GSVA enrichment scores of the pathways obtained for the DEGs with adjusted p-value <0.05. The enrichment score range is from -0.6 to 0.6.* *GBC, globose basal cell -like cells.*
